# Novel cell and tissue dynamics drive the unusual biology of the catch tentacle, an inducible organ of aggression found in the sea anemone *Metridium senile*

**DOI:** 10.64898/2026.04.13.718255

**Authors:** Rowan N. Lopez, Sarah E. Arnold, Kennedy Bolstad, Leslie S. Babonis

## Abstract

*Metridium senile* is a clonal anemone that engages in fighting interactions to defend its territory from non-clonemates utilizing an inducible fighting organ, the catch tentacle. Upon contact with a non-clonemate, the catch tentacle tip detaches onto the non-clonal individual, resulting in necrosis where the tip attaches to the non-clone. The incapacitating function of the catch tentacle is driven by a unique type of cnidocyte, the holotrich, which is not found elsewhere in *M. senile,* including the feeding tentacles from which catch tentacles develop. *Metridium farcimen,* the sister species to *M. senile*, never develops catch tentacles despite their close phylogenetic relationship, as exemplified by their ability to hybridize. Here, we compare the feeding tentacles of both species to the catch tentacles in *M. senile* to determine how catch tentacles achieve their unusual function. We found that the feeding tentacles of *M. senile* and *M. farcimen* house similar types of cnidocytes that develop from proliferative cells distributed throughout the tentacle. By contrast, catch tentacles house distinct cnidocyte types from feeding tentacles and restrict proliferative cells to the base of the tentacle. This suggests immature cnidocytes migrate from base to tip to replace lost cells after an aggressive interaction in the catch tentacle. Additionally, we observed two morphologically and chemically distinct types of holotrichs in the catch tentacles that appear to use different cues to induce firing. Together, our data suggests that the novelty of catch tentacle aggression is mediated by distinct cell and tissue dynamics.

## Introduction

Across the animal tree of life, organisms exhibit an array of mechanisms to protect themselves and their territory from other individuals. Sessile animals face extra pressures to maintain their territory as their limited mobility removes the option to inhabit a new territory. In anthozoan cnidarians (sea anemones, corals, tube anemones, and their relatives), some clonal species exhibit inducible organs of aggression which are used exclusively to compete for territory against other individuals [1]. These tissues are not present in every individual of a species, instead developing in lineages that must compete for territory with non-clonal conspecifics or other species [1]. These tissues are populated by a distinct type of cnidocyte (stinging cell) thought to be specialized for aggressive interactions [2]. In some scleractinian corals, sweeper tentacles are induced at the periphery of colonies to maintain an individual’s territory, causing lesions and necrosis on aggressor corals that grow too close to an individual [1,3,4]. In certain sea anemones, knob-like structures called acrorhagi inflate from a region of the body wall under the feeding tentacles and are used to make aggressive contact with other individuals. After contact is made, the tip detaches, causing necrosis at location of attachment on the victim anemone [5,6]. Catch tentacles are another inducible aggressive organ found in some species of sea anemones. Similar to acrorhagi and sweeper tentacles, upon contact with a non-clonal individual, the tip of the catch tentacles detaches leading to necrosis in the intruder [1]. In all these inducible organs of aggression, it is the cnidocytes, or stinging cells, which are responsible for the necrosis and tissue degeneration that results from these aggressive interactions.

Cnidocytes (stinging cells) are a cell type found only in cnidarians. Inside the cnidocyte is the cnidocyst, an organelle characterized by a pressurized capsule which houses an eversible, venom-filled tubule [7]. A sensory cone composed of stereovilli and a kinocilium sits on top of these cells, detecting chemicals or vibrations near the cell which allow the cnidocyte to determine prey movement and discharge its cnidocysts more effectively for prey capture discharge [8,9]. The venom filled tubule of the cnidocyst everts and releases its venom, an action which requires the cnidocyte to be replaced [10].

*Metridium senile,* the plumose sea anemone, is a species of sea anemone which can induce the development of catch tentacles from feeding tentacles [11,12]. During this transition, the suite of cnidocytes in the feeding tentacle changes, with a new type of cnidocyte dominating the tentacle, the holotrichous isorhiza (hereafter “holotrich”) [11,13,14]. Catch tentacles are not found on every *M. senile* individual as induction of this structure is tied to social context rather than being constitutive [11]. While *M. senile* can reproduce both sexually [15,16] and asexually through pedal laceration [17], it is the asexually derived individuals at the boundaries between competing colonies that tend to develop catch tentacles[11]. *Metridium senile* catch tentacles are capable of movement in their deflated form (Fig 1A). Aggressive behavior is initiated when the catch tentacle inflates and exhibits a searching motion through the water column until contact with another individual is established. At this point, the tentacle is inflated (Fig 1B) [11,18]. After making physical contact, *M. senile* detaches the tip of the catch tentacle onto the intruding anemone. As the aggressor pulls back, the catch tentacle tears (Fig 1C) [11] in an action similar to the “ectodermal peeling” described in the fights with acrorhagi in other anemones [6]. The victim’s impacted tissue undergoes necrosis and, in rare cases, death of the whole animal (Fig 1D-E) [11]. The holotrichs in the catch tentacle are responsible for this attachment and venom induced necrosis (Fig 1F), as necrosis does not occur when individuals interact with the feeding tentacles of an aggressor [11]. Catch tentacle induction is not unique to *M. senile.* At least 16 species of sea anemones have the ability to make catch tentacles, which are characterized by the presence of holotrichs in the tissue [1]. Despite their wide use in a variety of species, little is known about the development of holotrichs in these inducible organs.

**Figure 1:**
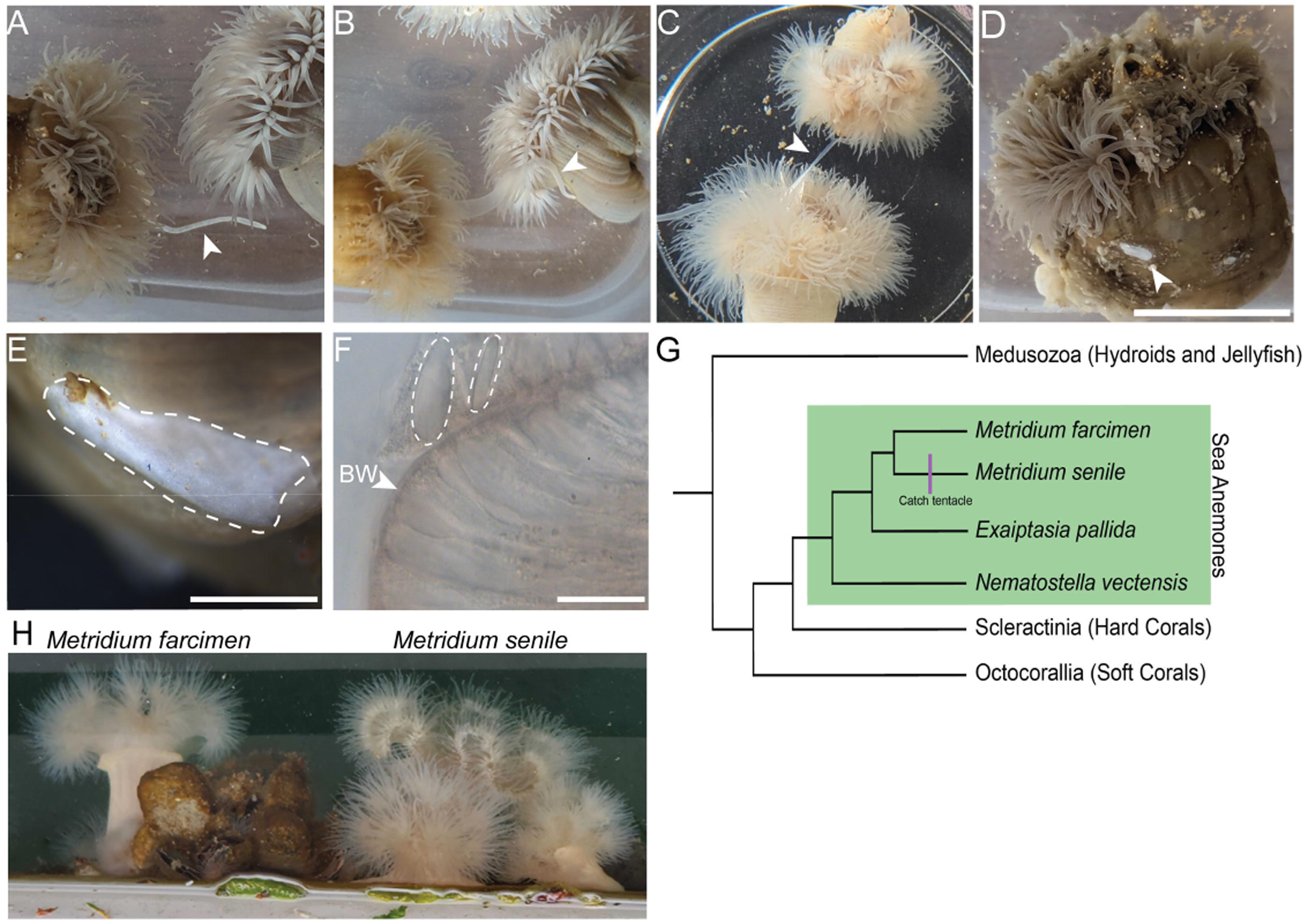
*Metridium senile* uses its catch tentacle for intraspecies aggression. **A.** *M. senile* individuals. One has an extended catch tentacle (arrowhead) at the start of searching behavior but prior to tentacle inflation **B.** Catch tentacle inflation and contact (arrowhead) with a nearby individual. **C.** Catch tentacle retraction: the attacker retracts the catch tentacle leaving the tip on the victim. Site of tip detachment shown by white arrowhead. **D.** Victim anemone following an attack. Catch tentacle tip (arrowhead) remains attached to the attacked individual. **E.** High magnification image of a catch tentacle tip still adhering to the body of a victim anemone, 24 hours after the fight. **F.** DIC image of discharged holotrichs (dotted white outlines) from a detached catch tentacle tip on the body wall (BW) of a victim anemone, 24 hours after a fight. **G.** Cladogram of cnidarians. Catch tentacles (hashmark) are only found in *M. senile*, not its sister taxon, *Metridium farcimen.* **H.** *M. farcimen* and multiple individuals of *M. senile* on a boat in the Friday Harbor public marina. Scale bar in D = 5cm and applies to A-D. Scale bar in E = 50 mm. Scale bar in F = 20 µm.

*Metridium senile* has become an important species for studies examining the development of catch tentacles [11,13,14,18,19]. Importantly, its sister species *Metridium farcimen,* the white-plumed sea anemone, does not have the ability to make catch tentacles, allowing for a comparison of tentacle cell dynamics in two closely related species (Fig 1G). *Metridium farcimen* is found across the Pacific from Kamchatka, Russia to at least the Gulf of California, having some overlap between the Atlantic and Pacific range of *M. senile* [20]. In the regions of overlapping distribution, *M. farcimen* and *M. senile* are often found close together (Fig 1H), with hybridization occurring frequently between the two species [21]. Despite experiencing similar environmental cues, *M. farcimen* only reproduces sexually [17] and will aggregate in groups of non-clones, whereas *M. senile* can reproduce sexually [15,16] or asexually by pedal laceration which results in clonal groups [17]. To understand how *M. senile* acquired the capacity to make catch tentacles, we compared the cell and tissue dynamics of the catch tentacles to the feeding tentacles in *M. senile*. To further determine if feeding tentacles in *M. senile* have a special capacity to become catch tentacles, we compared the feeding tentacles of *M. senile* to the feeding tentacles in *M. farcimen.* We show the distribution and types of cnidocytes in the feeding tentacles is similar in both species, and that holotrichs are located only in the catch tentacle of *M. senile.* To determine how cnidocytes are replaced in the different types of tentacles, we looked at the distribution of immature cnidocytes and proliferating cells. Given the different functions of the feeding tentacles and catch tentacles, we expected there to be a difference in the distribution across tentacle types. We further expected differences between the feeding tentacles across the two species as *M. senile* feeding tentacles can develop into catch tentacles. We show that immature cnidocytes are distributed throughout the tentacle regardless of type. Cell proliferation was regionalized only in the catch tentacles, with proliferative cells being present only at the base of the tentacle and absent at the tip, where aggressive interactions occur. Finally, we examine the chemical and morphological features of the two types of holotrichs in the catch tentacles. We describe key differences in apical sensory structures and chemical components, suggesting these two cell types play different roles in detecting the presence of non-clonemate and delivering venom.

## Methods

### *Metridium* culture and tissue collection

Sea anemones were collected from the public docks at the port of Friday Harbor on San Juan Island and Anacortes public marina in Washington State, USA. All *Metridium senile* were identified through the presence of catch tentacles. All animals were shipped to Ithaca, NY where they were maintained at 10□in 1X filtered seawater (Tropic Marin) on a 12-hour light cycle and fed freshly hatched *Artemia salina* nauplii once or twice per week. Prior to tissue collection, animals were starved for one week. To immobilize animals, menthol crystals were placed in 1X filtered seawater with the animals for up to 5 hours at 10□. Immobilization was determined by a lack of response after agitation with a pipette and allowed for full tentacle clipping at the base where the tentacle meets the oral disc for fixation and analysis of complete tentacles.

### Quantitative cnidocyte analysis

Tentacles were fixed for 90 seconds in 4% paraformaldehyde with 0.2% glutaraldehyde in phosphate-buffered saline with 0.1% Tween 20 (PTw) at 22□ (room temperature). Initial fixative was removed and tissues were further fixed in 4% paraformaldehyde in PTw at 4□for an additional 12-18 hours. Fixed tentacles were washed three times in PTw to remove excess fixative and stored at 4□in clean PTw before processing. To determine the proportion of each type of cnidocyte in the tentacles, an individual tentacle was placed on a glass slide with 80% glycerol diluted in PTw. A coverslip was gently lowered onto the tentacle and pressed down until there was no visible intact tissue. To make a permanent prep of the tissues, the coverslips were sealed onto the slide with nail polish. Images were taken covering the entire squashed tissue area using the tiling function on a Zeiss LSM 900 upright confocal microscope. FIJI [22] was used to overlay a grid with areas of 20,000 microns^2^ and a number was assigned to every box in the grid. Numbers were then randomly selected utilizing the site https://www.random.org/ to identify 20 grids for analysis of cnidocyte distribution. Any grids that lacked cnidocytes were discarded and random numbers were chosen again to ensure there were 20 replicate grids containing cnidocytes. Lengths and widths were measured for each cnidocyte that was in focus in the grid. Counting was repeated for three feeding tentacles from each of three individuals for both species (N = 9 tentacles for each species). For the catch tentacles, five tentacles were analyzed, one from each of five individuals. Catch tentacles were first cut into three sections (tip, middle, and base) before squashing each section independently.

### Cnidocyte staining

To visualize mature cnidocytes in intact tissues, tentacles were fixed for 90 seconds in 4% paraformaldehyde with 0.2% glutaraldehyde in phosphate-buffered saline with 0.1% Tween 20 (PTw) at 22□ and refixed for 1 hour in 4% paraformaldehyde and 10mM EDTA in 1X Tris Buffer at 4□ (following the protocol of Szczepanek et al., 2002) [23]. Following fixation, tissues were washed 3 times in PTw and cnidocytes were labeled by incubation in 143 µM DAPI diluted in PTw for 30 minutes at 22□. Excess DAPI was removed with 3 washes in PTw, and tissues were mounted in 80% glycerol on slides coated in Rain-X (Illinois Tool Works, USA) for imaging.

### Immunohistochemistry

To characterize immature cnidocytes and sensory cone morphology, tissues were fixed as for squash preps, with the post fix (4% paraformaldehyde in PTw) for 1 hour at 4□. Tissues were then washed 3 times in PTw to remove excess fixative and incubated for one hour in PBS with 0.2% Triton X-100 and 0.1% BSA (PBT). This solution was removed and tissues were incubated in 5% normal goat serum in PBT for 1 hour at 22□. Tissues were then incubated at 4□ overnight in primary antibody diluted in 5% normal goat serum in PBT. A *Nematostella vectensis* specific antibody against minicollagen 4 [24] was used 1/500 to label immature cnidocytes in both species of *Metridium.* A commercially available antibody recognizing acetylated tubulin (Sigma T6743) was used 1/200 to label cilia, and an antibody recognizing phosphorylated histone H3 (hereafter: PH3) [25] was used 1/50 to detect proliferating cells. Primary antibody was removed with 5 washes in PBS with 0.2% TritonX and tissues were then incubated overnight at 4□ in goat-anti-rabbit secondary antibody for minicollagen 4 and for one hour at 22□ in goat-anti-rabbit (for PH3) and goat-anti-mouse (for acetylated tubulin) secondary antibodies diluted at 1/500 in PBS with 0.2% Triton X. For acetylated tubulin labeling, tissues were also counterstained with phalloidin (Invitrogen A12379) diluted 1/200 in PBS with 0.2% TritonX to visualize F-actin. As an alternative way of labeling proliferating cells, tentacles were also analyzed using a Click-iT® EdU Imaging Kit (Invitrogen C10340). Prior to fixation, tissues were incubated for 30 minutes in 10 1X filtered seawater at room temperature (22□) in 100µM EdU. After incubation, 100 µL of 7.14% MgCl_2_ were added to the test tubes containing EdU for 10 additional minutes. Tentacles were fixed as described above for other immunohistochemistry fixations. To visualize proliferating cells with EdU, tissues were washed in PTw for 5 minutes at 22□ before being washed for 20 minutes at 22□ in PBS with 0.5% Triton X-100. Tissues were then incubated in EdU Click-It reaction cocktail (prepared as described by the manufacturers) for 30 minutes at 22□. Tentacles were washed in several changes of PTw for 30 minutes to remove reaction cocktail. Nuclei were counterstained with a 30-minute incubation in 1.43 µM DAPI at 22□. Tissues were mounted in 80% glycerol and imaged on a confocal microscope.

## Results

### *Metridium farcimen* feeding tentacles are indistinguishable from *M. senile* feeding tentacles

Despite previous work on catch tentacle development in *Metridium senile* [11,13,14,18,19], little is known about how the species evolved the capability to induce these organs of aggression due to the limited knowledge of *Metridium farcimen.* Given that *M. farcimen* does not induce catch tentacles, we expected the feeding tentacles of *M. farcimen* to differ from the tentacles of *M. senile,* with the catch tentacles and feeding tentacles of *M. senile* being most similar. To determine potential differences between *M. senile* and *M. farcimen*, we analyzed the cnidocyte composition of the feeding tentacles of the two species (Fig 2A, B). Three types of cnidocytes have been described in the feeding tentacles of *M. senile* [13,14], two nematocytes: basitrichous isorhizas (also called microbasic b-mastigophores; hereafter “basitrichs”) and microbasic p-amastigophores (hereafter “amastigophores”), and spirocytes. These cnidocytes were also found in the feeding tentacles of *M. farcimen* in similar proportions to the feeding tentacles of *M. senile* (Fig 2C). Spirocytes were the dominant type of cnidocyte in the feeding tentacles, making up approximately 79% of the cnidocytes in *M. farcimen* feeding tentacles and 73% in *M. senile,* supporting a previous observation that this type of cnidocyte was most dominant in *M. senile* feeding tentacles [14]. Amastigophores were the second most common cnidocyte type in the feeding tentacles, making up 11% of the cnidocytes in *M. farcimen* and 18% in *M. senile*.

**Figure 2:**
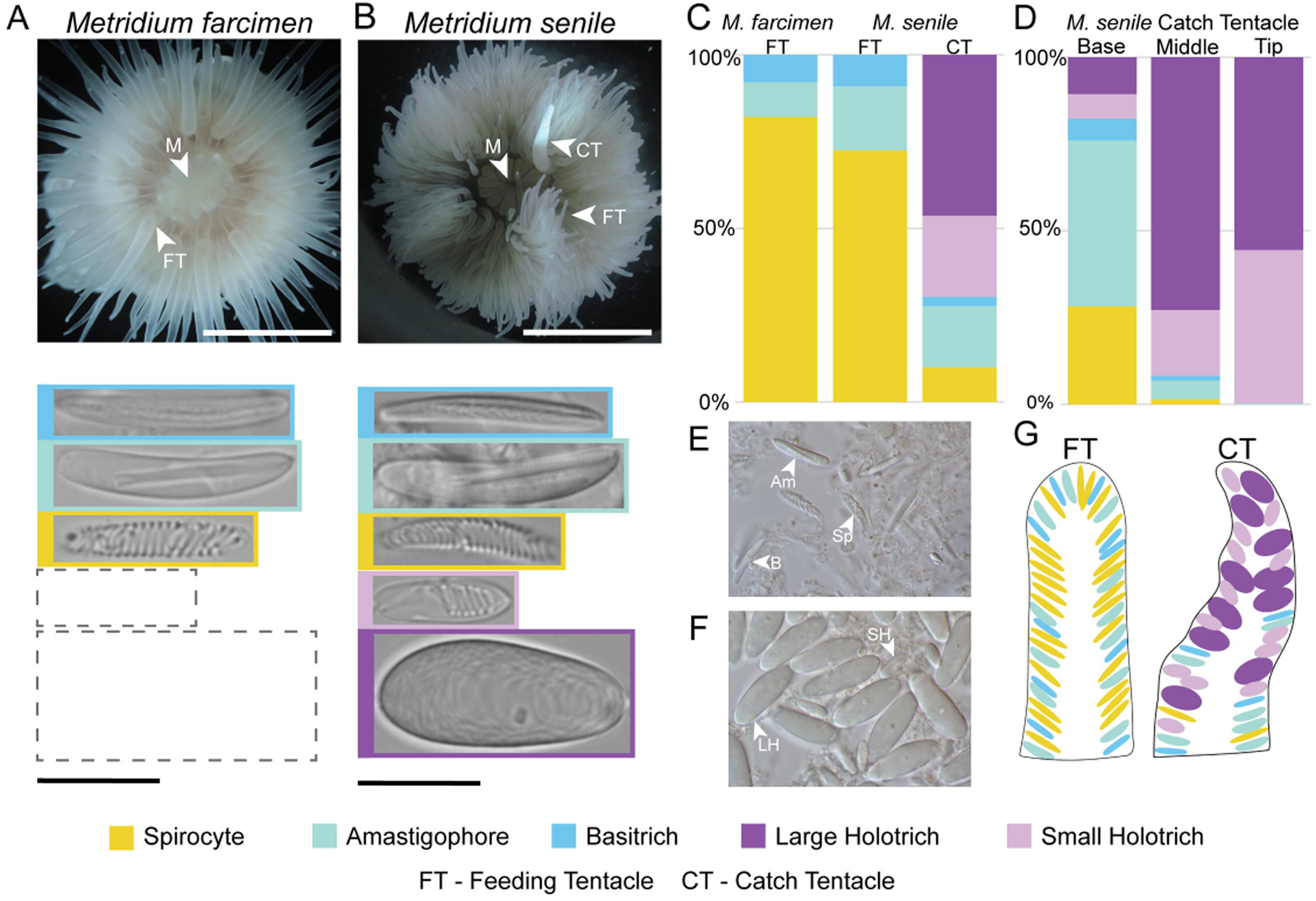
Characterization of cnidocytes in *Metridium farcimen* and *Metridium senile* tentacles. **A-B.** Oral view of adult animals indicating feeding tentacles (FT), catch tentacle (CT), and mouth (M). Cnidocytes found in the tentacles of *M. farcimen* (left) and *M. senile* (right) are shown underneath the animal. Both species have basitrichs (blue), amastigophores (green), and spirocytes (yellow). Holotrichs (two types, indicated by light and dark purple) are only found in *M. senile.* **C.** Proportion of each cnidocyte type across tentacles. Feeding tentacles are indistinguishable between species; holotrichs are only found in the catch tentacles. **D** Proportion of cnidocytes from the base to the tip of the catch tentacles. Cnidocytes shared with the feeding tentacles are regionalized towards the base of the catch tentacle. **E.** Representative DIC image of dissociated cnidocytes from the feeding tentacle of *M. senile* **F.** Representative DIC image of dissociated cnidocytes from the catch tentacle of *M. senile* **G.** Schematic of cnidocyte distribution in different tentacle types. Feeding tentacle cnidocytes are smaller and more elongate than catch tentacle cnidocytes. Scale bar for *M. farcimen* in A = 1 cm. Scale bar for *M. senile* in A = 2 cm. Scale bar for cnidocytes in A and B = 10 µm.

Basitrichs were the least common cnidocyte, making up 10% and 9% of the total cnidocyte proportion in *M. farcimen* and *M. senile* respectively. These proportions were similar to what has been previously described for *M. senile* feeding tentacles, where amastigophores and basitrichs were reported to be at roughly the same abundance [14]. we observed no distinct size classes when measuring the cnidocytes from the feeding tentacles of either species; rather, lengths of the cnidocytes were continuously distributed for spirocytes, amastigophores, and basitrichs (Supplemental Table 1). This was unexpected as previous works have shown distinct size differences in the cnidocytes of *Nematostella* [26] and previous studies on *Metridium* grouped cnidocytes on different sizes [14]. Size of the anemone or tentacle did not change the range of cnidocyte sizes observed, consistent with previous reports on *Metridium* tentacle cnidocyte scaling which showed similar sizes across differing body sizes [27].

### Large and small holotrichs are found in different proportions throughout the catch tentacle

No holotrichs were found in *M. farcimen*. In *M. senile,* we found two types of holotrichs which were abundant in the catch tentacles (Fig 2C) as previously reported [13,14]. The size of the holotrichs were similar to those reported by Östman et al. (2010) [14] and were not present in the feeding tentacles, as expected based on previous work [11]. In addition to size, the two types of holotrichs had clear morphological differences: the coiled tubule of the small holotrich was clearly visible through the capsule and was positioned towards the apical side of the elongate and small capsule whereas the coiling of the large holotrich tubule was less ordered and could be seen throughout the rounder and larger capsule (Fig 2B) [13]. Large holotrichs were the most prevalent cnidocyte in the catch tentacles, making up 46% of the tentacle cnidocytes, followed by the small holotrichs which made up 23% (Fig 2C). Initial observations suggested to me that unlike the feeding tentacles, cnidocytes were regionalized in the catch tentacle (Fig 2D). To determine where these cells are found in the catch tentacles, we cut the tentacles into three sections, a tip region, a middle region and a basal region before squashing and imaging (Fig E, F). At the tip of the tentacle, large and small holotrichs compromise nearly 100% of the cnidocytes (56% and 44%, respectively) (Fig 2D). Large holotrichs were more prevalent than small holotrichs in the middle section (73% and 19% respectively) but these proportions even out at the base where large holotrichs make up 11% of the cnidocytes and small holotrichs 7% (Fig 2D). These differences suggest small and large holotrichs are being used differently in the aggressive behavior of the catch tentacles. Other cnidocytes were present in the catch tentacles (Fig 2C) but were restricted to the base and middle section of the catch tentacle (Fig 2D). At the base, amastigophores were the most dominant cnidocyte shared between the feeding tentacles and catch tentacles (48%), followed by spirocytes (28%) and basitrichs (6%) (Fig 2D). The proportion of feeding tentacle cnidocyte decreases in the mid-section, with amastigophores being the most prevalent (5%) and spirocytes and basitrichs being present at similar proportions in this region (both 1%) (Fig 2D). The occasional amastigophore was found at the tip of the catch tentacle (Fig 2D). In summary, cnidocytes in the feeding tentacles were evenly scattered throughout the tissue whereas the dominant cnidocytes in the catch tentacle were regionalized toward the tip (Fig 2G).

### Cnidocyte are replaced from the base of the catch tentacle

After a cnidocyte fires, the cell dies and must be replaced by a progenitor cell [10]. Given the distinct cnidocytes of the catch tentacle, we wondered if the catch tentacles housed a distinct population of progenitor cells. To determine how cnidocytes are replaced in the tentacles, we examined the distribution of proliferative (progenitor) cells and immature cnidocytes using immunohistochemistry. Based on previous studies of the burrowing sea anemone, *Nematostella vectensis,* we expected proliferating cells to be present throughout the tentacles except for the tentacle tips [28], with immature cnidocytes present throughout the entire tentacle [24]. Using an antibody directed against a cnidocyte-specific protein (minicollagen), we found intracellular structures in feeding tentacles that had either a U or J shape (Fig 3 A-F). The structures were found in both *M. farcimen* and *M. senile* feeding tentacles and were similar to what has previously been described for developing/immature cnidocytes in *M. senile* [14]. When we overlaid the minicollagen4 labeling with DIC, it was clear the labeled structures were developing cnidocysts. This pattern of protein localization throughout the tentacle was also seen in the catch tentacles of *M. senile* (Fig 3G-I) though the shapes of the immature cnidocytes were very different from those seen in the feeding tentacles (e.g., Fig 3G insets). The unique shapes found in the catch tentacles corresponded to developing holotrichs, showing that minicollagen4 antibody labels holotrichs in addition to the known labeling of nematocytes and spirocytes in *Nematostella* [24].

**Figure 3:**
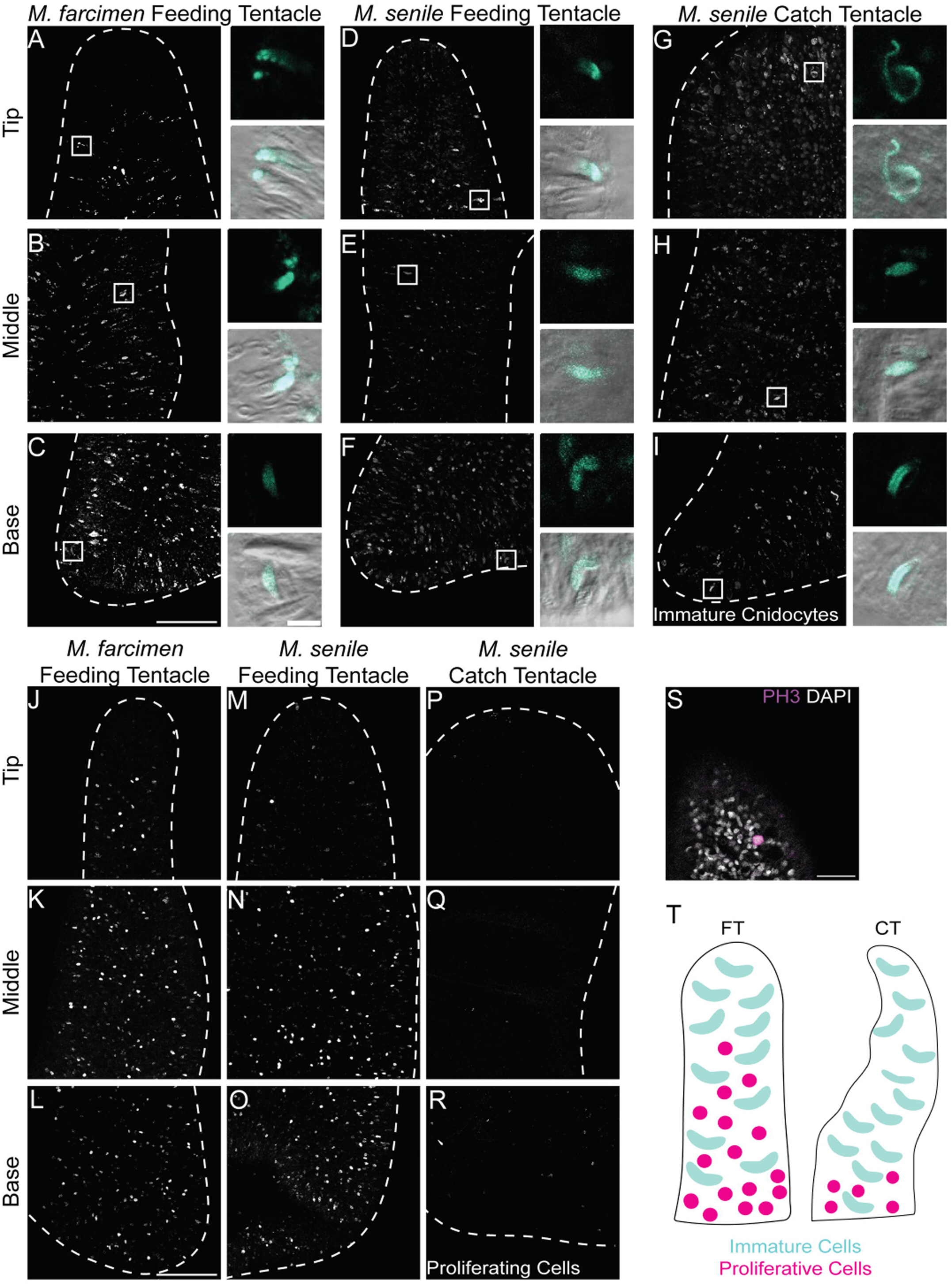
Development of cnidocytes across tentacle types. **A.-I.** Max projection of immature cnidocytes (white) labeled with anti-minicollagen 4 antibody. Tentacles outlined in dotted white line. Insets: Single optical sections of immature cnidocytes (cyan) overlaid with DIC. Insets are from regions outlined by white boxes in max projection images. **J-R.** Max projection of proliferative cells (white) labeled with anti-PH3 antibody. Scale bar in bottom left applies to all max projections (50 µm). **S.** High magnification image of a proliferative cell (magenta) counterstained with DAPI, which labels all nuclei (white). **T.** Schematic of immature cnidocytes and proliferating cells in *M. senile* and *M. farcimen* tentacles. Feeding tentacle (FT) and catch tentacle (CT). Scale bar in C (max projections) = 100 µm and applies to max projections in A-I. Scale bar in C (inset) = 10 µm and applies to insets in A-I. Scale bar in S = 20 µm.

To determine where the developing cnidocytes were coming from, we labeled tentacles with an antibody directed against phosphohistone H3. This protein is phosphorylated in cells during M phase, and its localization has been shown to be a reliable marker of proliferating cells *N. vectensis* [29]. We found that proliferating cells were not restricted to a particular region of the feeding tentacles in *M. farcimen* or *M. senile*; instead, they were scattered throughout the ectoderm of the entire tentacle (Fig 3J-O) as seen in *N. vectensis* [28,30]. In contrast with the feeding tentacles, proliferating cells in the catch tentacles were restricted largely to the base of this tissue, with the abundance of proliferating cells decreasing from base to tip (Fig 3 P-R).

### Large holotrichs are more similar to spirocytes than previously expected

To visualize the location of mature cnidocytes in whole tentacles, we labeled tentacles with 143µm DAPI (Fig 4A). When introduced in high concentrations, DAPI binds to the poly-γ-glutamate found inside the pressurized capsule of mature nematocytes [23]. Spirocytes do not label with 143µm DAPI and as such, we expected to see only nematocytes labeled with DAPI across the tentacles. Mature, DAPI-labeled nematocytes were found from base to tip for both feeding tentacles and catch tentacles (Fig 4A), as expected from our analysis of cnidocyte distribution (Fig 2). Surprisingly, large holotrichs, one of the dominant nematocytes in the catch tentacles, did not label with DAPI though the small holotrichs did (Fig 4B). As poly-γ-glutamate is involved in pressurizing the nematocyte for firing, this suggests that the two types of holotrichs may use different polymers to create the osmotic pressure necessary for firing [23].

**Figure 4:**
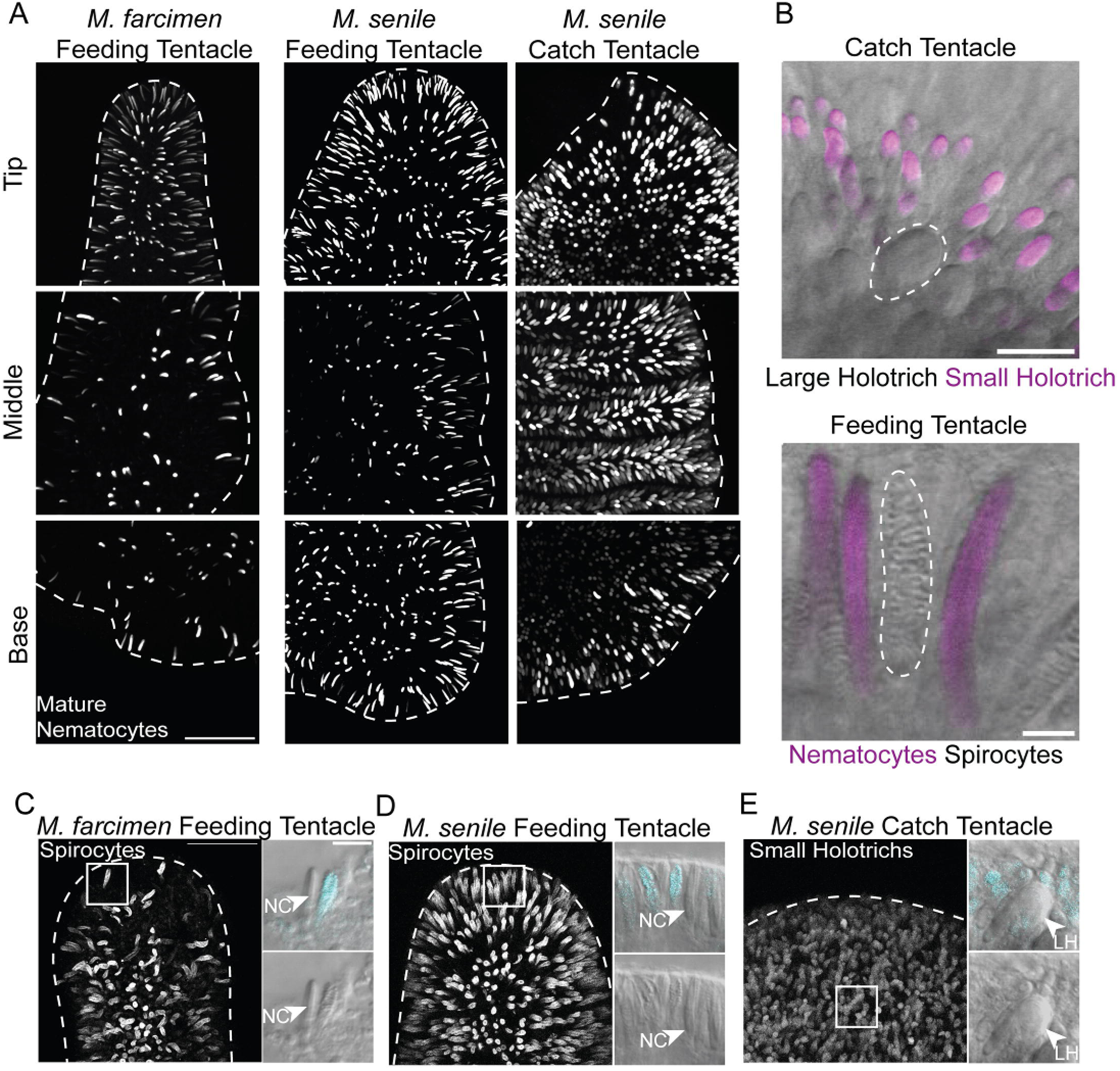
Large holotrichs look more like spirocytes than nematocytes with high concentration DAPI. **A.** Mature nematocytes labeled with 143 µM DAPI (white) across tentacles. Max projections are shown with tentacles outlined in white dotted lines. **B.** DIC images of cnidocytes in situ showing DAPI labeling (magenta) is absent from large holotrichs and spirocytes. Large holotrichs and spirocytes outlined in white. **C-E.** Alexa fluor labeling of spirocytes and holotrichs. Left, max projection of tentacle tip overview. Tentacles outlined in white dotted line. Boxes indicate location of insets. Insets: high magnification DIC images showing labeling (cyan) is restricted to spirocytes and small holotrichs. Other nematocytes (NC), including the large holotrichs (LH), (arrowheads) are not labeled. Scale bar in A = 100 µm. Scale bar in B (holotrichs) = 20 µm. Scale bar in B (feeding tentacles) = 5 µm. Scale bar in C (max projection) = 50 µm and applies to max projections in C, D and E. Scale bar in C (inset) = 10 µm and applies to insets in C, D and E.

These results also suggest the firing mechanism used by large holotrichs may be more similar to spirocytes than to small holotrichs. To further probe the relationship between large holotrichs and spirocytes, we developed a reliable method for labeling spirocytes using an Alexa Fluor azide (a component of the EdU kit described in Methods). We show abundant spirocytes in the tips of the feeding tentacles of both species (Fig 4C, D), supporting our analysis of spirocyte distribution (Fig 2). Nematocytes in the feeding tentacles of both species were unlabeled, allowing for optimal visualization of the spirocytes in these tissues. Interestingly, in the catch tentacles, large holotrichs were not labeled by this reagent while small holotrichs were (Fig 4E). we are unsure as to what Alexa Flmy azide 647 binds to in these cells; however, the Click-It chemistry relies on the presence of copper to catalyze the azide reaction suggesting maybe spirocytes and small holotrichs sequester copper. Alternatively, this azide labeling could be related to intracapsular pH as previous work has shown that unfired spirocytes can be labeled with acidic dyes [31]. These results suggest further investigation of the intracapsular makeup of holotrichs is warranted.

### Large holotrichs and spirocytes lack apical sensory cones

Previous work on *Metridium senile* described the presence of a sensory structure composed of a ring of microvilli surrounding a cilium on the apical regions of the feeding tentacle nematocytes [13]. To visualize the apical sensory cone morphology of the cnidocytes in the tentacles, we labeled tentacles with phalloidin (to detect filamentous actin in microvilli) and an antibody directed against acetylated tubulin (to detect cilia) in *M. senile* (Fig 5). The catch tentacles exhibited prominent cones (Fig 5A) and abundant actin rings (Fig 5B). In contrast with what has been described previously, we show that these apical actin rings are associated with the large holotrichs but that there is no cilium associated with this structure (Fig 5C-E). By contrast, the prominent cones with long cilia were associated with the small holotrichs (Fig 5F-I), as described previously [13]. The sensory cones were smaller and sparser on the feeding tentacles of *M. senile* (Fig 5J-K). The pattern of sparse sensory cones was also observed on the feeding tentacles of *M. farcimen* (data not shown). This was expected due to the high proportion of spirocytes in the feeding tentacles, which lack apical sensory cones and are instead topped by a small ring of actin-based microvilli (Fig 5K-N) [31]. As expected, nematocytes in the feeding tentacle had an apical sensory cone with a cilium (Fig 5O) similar to the cones of the small holotrichs, though the cones of the feeding tentacle nematocytes were smaller than those of the small holotrichs. The similarity in the apical morphology of spirocytes and large holotrichs further supports a close relationship between these two cell types.

**Figure 5:**
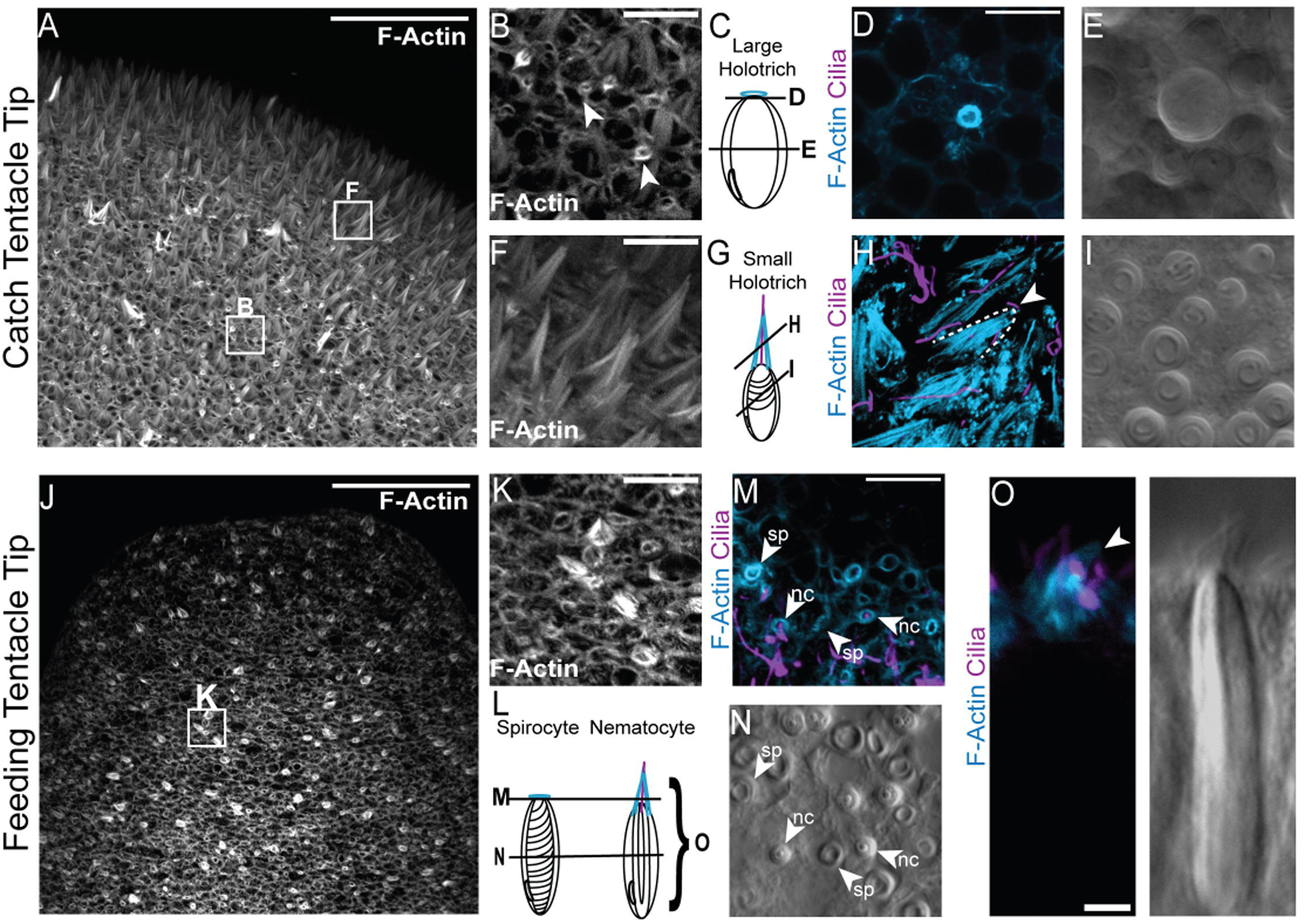
Apical sensory cone morphology varies across cnidocyte types. **A.** Max projection of F-actin labeling in the catch tentacle tip. **B.** High magnification region of catch tentacle indicated by the box in A. Actin rings are present throughout the tentacle (white arrowheads). Scale bar is 10 µm. **C.** Schematic of a large holotrich with an apical actin ring. Solid black lines correspond to image regions of panels D and E. **D.** Optical section of the apical structure of a large holotrich showing an actin ring (blue) with no cilium. **E.** DIC image of a single optical section through the capsule of a large holotrich. **F.** High magnification region of the catch tentacle tip indicated by the box in A showing tall cones (white). **G** Schematic of a small holotrich with tall apical cone. Solid black lines correspond to image regions shown in panels H and I. **H.** Max projection of apical structure of several small holotrichs at an oblique angle. Sensory cone (blue) has an associated cilium (purple). Cone outlined by dotted line with arrowhead showing associated cilium. **I.** DIC image of a single optical section through a field of small holotrichs. **J.** Max projection of F-actin (white) labeling of *M.senile* feeding tentacle tip. **K.** High magnification image of a region of the feeding tentacle corresponding to the box in J. Actin rings and short sensory cones (both white) are found throughout the tentacle. **L.** Schematic showing the apical actin ring on a spirocyte and a short apical sensory cone on a nematocyte. Solid black lines correspond to image regions shown in M and N, bracket indicates a side view shown in O. **M**. Max projection of spirocytes and nematocytes showing apical structures. Spirocytes (SP) have a ring of actin (blue). Nematocytes (NC) have an actin cone (blue) with cilium (purple). **N.** DIC cross section of spirocytes and nematocytes in a *Metridium senile* feeding tentacle. **O.** An apical sensory cone on an amastigophore nematocyte (arrowhead). Left, max projection of a short sensory cone (blue) with cilium (purple). Right, same image with DIC. Scale bar in A, J = 50 µm. Scale bar in D, F, K, M = 10 µm and applies to D, E, F, H and I, K, M and N. Scale bar in O = 3 µm.

## Discussion

### Similarities between Metridium farcimen and Metridium senile

Previous taxonomic work in cnidarians has included the characterization of cnidocytes, showing that the general types of cnidocytes within a genus remain the same while sizes have some variation [32]. While comparisons between *M. senile* and *M. farcimen* have focused on another aggressive tissue, the acontia [27], in this study we showed the feeding tentacles of *Metridium senile* and *Metridium farcimen* are similar in the types of cnidocytes they house (Fig 2, Supplemental Table 1) and their mode of cnidocyte replacement (Fig 3). While composition of cnidocytes in an animal is not enough to determine species, with most anthozoans sharing the same general suite of cnidocytes (spirocytes, microbasic p-amastigophores, basitrichs and/or microbasic b-mastigophores) [32–34], the holotrichs stand out in *M. senile* as a cnidocyte type which correlates to a specific, inducible, tissue with a specialized function in aggression. This work supports the findings of Purcell 1977 which showed that catch tentacle development involves the production of holotrichs and are absent in the feeding tentacles of *M. senile* [11]. While the commonalities of the feeding tentacles are not surprising as the two species can hybridize, the presence of holotrichs only in the catch tentacles reemphasizes the novelty of catch tentacles in *Metridium senile*.

### Cnidocyte replacement reflects tentacle function

Regarding cnidocyte replacement, we saw similar patterns across the three tentacles wherein immature cnidocytes were located throughout the tentacle (Figure 3). The location of immature cells in *Metridium* tentacles displays a different pattern than those found in another anemone which develops catch tentacles, *Haliplanella (Diadumene) luciae.* In *Haliplanella,* immature cnidocytes are present but only occasionally in the feeding tentacles and early developing stages of catch tentacles, with a complete absence in the mature catch tentacle [12], a stark difference to the clear prevalence in *Metridium* (Fig 3A). Proliferative cells were least prevalent at the tip of the feeding tentacles and were completely absent from the catch tentacle. Previous work on the development of catch tentacles in *M. senile* and *H. luciae,* has shown that during the development of a catch tentacle from a feeding tentacle, the percentage of feeding tentacle cnidocytes decreases and this is followed by the appearance of holotrichs [11,12]. Our results suggest that developing holotrichs differentiate as they travel up to the tip of the catch tentacle as immature cnidocytes were seen throughout the entire tentacle while proliferative cells were restricted to the base. Additionally, the more prevalent curvature of cells at the basal regions suggests that the cells at the base are younger than those found in the tip. The migration of cells was similarly suggested by Watson and Mariscal (1983) in their study of *H. luciae* catch tentacles to describe the surge of immature cnidocytes seen in the middle stages of catch tentacle development [12].

Given that the catch tentacle tip is “sacrificed” for the aggressive interaction, we suggest that the placement of proliferative cells at the base may be a key to maintaining the ability to replace the tip after it is lost. The presence of immature cnidocytes at the tip of the catch tentacles suggests that the tissues collected were either still developing while they were engaging in the searching behavior or were already replacing fired cells. Alternatively, the presence of immature cnidocytes in the catch tentacle tip may suggest that catch tentacles fill their tips with immature cnidocytes that only continue developing once tip detachment occurs. This could prolong the venom exposure to the victim anemone after the fight and worsen the necrotic effect.

### Large holotrichs share chemical and morphological properties with spirocytes

Holotrichs are a type of cnidocyte found across the Cnidarian phylum. In *Hydra*, holotrichs are known to be penetrating cells and are the size of the small holotrichs in *Metridium* [35]. In the parasitic cnidarian, *Polypodium hydriforme,* four morphologically different types of holotrichs have been described, some of which are in the proximal tentacle and mouth which aid as penetrants in prey capture and others which are localized at the tips of sensory tentacles, which never fired when prey was presented [36]. In the anthozoans, holotrichs are present in the three inducible aggressive organs (sweeper tentacles, acrorhagi and catch tentacles) [1,4]. Studies on *Metridium senile* have described two types of holotrichs (holotrichs and atrichs in older literature) with differences in tubule coiling and capsule morphology [13] though the correlation to function is still unknown. Furthering this research, we determined key firing morphological and chemical differences between these cnidocytes. Small holotrichs most closely resembled other nematocytes in the feeding tentacle with the cell labeling with high concentration DAPI staining as in other nematocytes [23] (Fig 4). We further identified an apical cone associated with these small holotrichs (Fig 5), similar to the cones seen in other nematocytes in *Metridium* [13] and across anthozoans (reviewed by Babonis, 2025) [37]. When we examined the large holotrichs, we found no staining by high concentration DAPI, off target labeling with Alexa fluorophore and a lack of a sensory cone, instead showing an actin ring (Figs 4, 5). Previous observations on the apical structure of *M. senile* large holotrichs have shown a cilium associated with the apical section of the cell [13]. Our observations of the large holotrichs were instead most similar to the observations we made in the spirocytes of *M. senile* and *M. farcimen,* including the presence of a ring of short microvilli on the apical region of the cells as previously reported for spirocytes [31]. In their study of spirocyte apical morphology, Mariscal et al. (1976) suggested that the two types of microvilli of the spirocyte apical rings had individual mechanical and chemical sensory roles which may both be involved in the overall firing of the cell [31]. Given the poking behavior of *M. senile* before a fight and the chemical similarities, we suggest that the microvilli of the large holotrichs work in a similar fashion to the spirocytes. This could allow for greater sensing to determine the location of other anemones and aid in determining when a new non-clonal individual inhabits the territory.

Cnidarian allorecognition systems, the systems which determine an individual’s own tissues versus those from a different individual, have been studied most extensively in the colonial hydroid *Hydractinia* [38–42] (Reviewed by Rosengarten and Nicotra, 2011 and Nicotra, 2022) [43,44]. In these colonial animals, two allorecognition genes (Alr 1 and Alr 2) determine the relatedness between individuals through protein binding on the cell surface of their stolons, tissues that anchor the animals on substrate and can also be used to fight other colonies (reviewed by Nicotra, 2022) [44]. Similar to the catch tentacles, contact can result in nematocyte firing between non-clones and lead to necrosis [41] and these tissues can be induced when a new colony is introduced [40]. While the *Metridium* allorecognition system has not been studied, it is likely that the actual method for recognizing clonal individuals lies in the catch tentacle, with membrane proteins that may play similar roles as Alr 1 and Alr 2. The allorecognition system of *Metridium* then likely involves a chemical cue which prevents the firing of cnidocytes before and during the recognition stage of fighting. This type of control of cnidocyte firing has been observed in the feeding tentacles of *Hydra*, where well-fed animals will not continue to discharge cnidocytes after they have reached satiety [45]. It is possible that after a cue is received and cnidocyte firing is no longer inhibited that the holotrichs of *M. senile* can fire to attach the catch tentacle tip onto the victim anemone and begin releasing venom.

The role of the two types of holotrichs during tip adhesion is still uncertain. In nematocytes, the osmotic pressure behind cnidocyte firing is generated by poly-γ-glutamate right before the maturation of the cell, causing the capsule of the cnidocysts to swell and straighten [46]. High concentration DAPI binds to this polymer, resulting in the labeling of mature capsules [23]; DAPI similarly labeled the small holotrichs, suggesting that this polymer too is required for small holotrich firing. Given the rapid firing of nematocytes [47] and the use of holotrichs in other species for prey capture and defense [35,36] we suggest that small holotrichs are responsible for the main attachment of the tentacle tip on the victim anemone. This is not to say that the large holotrichs do not contribute to tentacle tip adhesion. In anthozoan feeding tentacles, spirocytes and nematocytes are thought to work together to capture prey items, with the spirocytes helping to entangle prey while the nematocyte delivers the venom [31]. We suggest a similar co-reliance occurs in the catch tentacles of *M. senile,* where one type of holotrich aids in attaching to the victim anemone while the other is more specialized to the venom delivery. Regarding their development, Watson and Mariscal’s study (1983) showed that large holotrichs appear once a suite of small holotrichs is already established in the developing catch tentacles of *Haliplanella luciae* [12]. This could suggest that small holotrichs are used for detection of the allorecognition cue and large holotrichs participate later to deliver the venom. Alternatively, it could simply reflect that large holotrichs take a longer time to reach maturity.

While catch tentacles stand out in *Metridium* as a novel, inducible fighting structure, the feeding tentacles in *M. senile* and *M. farcimen* were indistinguishable. Given the various similarities between the feeding tentacles, namely the process for replacing cells and the composition of cnidocytes they contain, a future investigation of the genes expressed in the feeding tentacles and catch tentacles of *Metridium senile* to determine what genes are responsible for catch tentacle development and whether they are shared with *M. farcimen.* Given the stark differences between the two types of holotrichs, it is possible that novel genes may be controlling the development of these cell types. Alternatively, the genes behind the development of the cnidocytes in the feeding tentacles may have been repurposed to give rise to the two distinct types of holotrichs in the catch tentacles. The tentacles of *Metridium senile* and *Metridium farcimen* provide a system in which to study the development and origin of new cell types through the repurposing of conserved genes and the origin of novel genes.

## Acknowledgements

This work was supported by the National Institutes of Health (R35GM147253-01 to LSB) and institutional funds from Cornell University (the Hunter Rawlings the Third Presidential Research Scholars to RNL) and funds from the Society of Developmental Biology (Choose Development! Program (R25-HD105600) to RNL). Quantitative image analysis was performed with Imaris Software (Oxford Instruments, UK) at the Cornell Institute of Biotechnology’s BRC Imaging Facility (RRID:SCR_021741).

## Figure Captions

**Supplemental Table 1:** Average size and standard deviation (µm) of unfired cnidocytes in the feeding tentacles and catch tentacles of *M. senile* and *M. farcimen*

## Notes

### Competing Interest Statement

The authors have declared no competing interest.

